# Agent-based Modeling of Asian Corn Borer Resistance to BT Corn

**DOI:** 10.1101/795724

**Authors:** Arian J. Jacildo, Jomar F. Rabajante, Edwin P. Alcantara

## Abstract

We present ABM-IRM, an agent-based modeling approach to insect resistance management. ABM-IRM is one of the agent-based models that we have developed to simulate the insect resistance of Asian Corn Borer (ACB) to BT corn. The model was implemented using NetLogo, an agent-based programming environment (Wilensky, 1999). We created our model using simple rules to find emergent patterns (Wilensky and Rand, 2015) based on refuge types and pyramid BT events that may aid in controlling the resistance of ACB to BT Corn. Following how the Overview, Design concepts, and Details (ODD) protocol was used in presenting a related ABM paper as guide (Anderson and Dragićević, 2015), the following sections are organized as follows: Section 2 covers the typologies of the agents, Section 3 discusses the process overview, and Section 4 highlights our results and emergent patterns.

## 2. Agents

### 2.1 Corn Field Agents

We modeled the corn field as a 2D grid of non-moving agents that we call patches in NetLogo. Green colored patches represent areas planted with BT corn and yellow patches are refuge areas. We investigated three types of refuge areas: “Circle”, “RandomSeeds” and “Strips”. Examples of these refuge types with varying proportion with respect to the total corn field area are shown in Figure 1. Circle and strip refuge types are explicitly setup in the field, whereas random seeds are incorporated in the distribution corn plant seeds.

**Figure 1.**
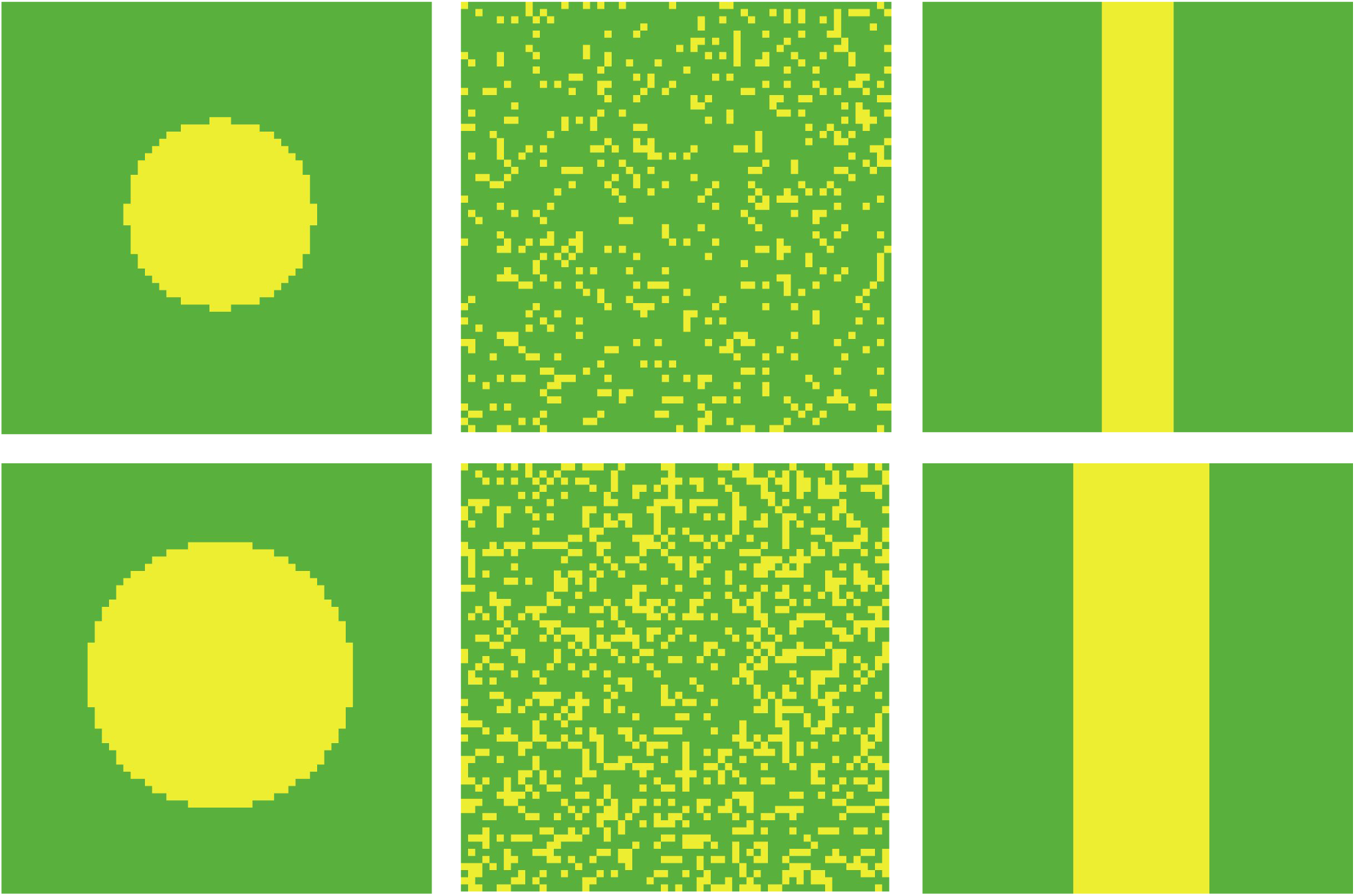
Varying refuge types, from left to right: circle, random seeds and strips. Sizes of refuge areas (yellow patches) in the first row and second row correspond to 15% and 30% of the total area, respectively.

### 2.2. ACB agents

We modeled ACB as mobile agents, or turtles they are called NetLogo. The simulation can be seen as the interaction among turtle agents over a period of time. We used the published life tables (Hussein et al. 1999) and life cycles (Lit et al., 1999) as assumptions to the properties of ACB agents. Table 1 shows the adapted assumptions based from cited literatures.

**Table 1.** Life table of ACB agents and calibrated mating parameters with their default values, definitions and assumptions.

| Parameter | Default Value | Definitions/Assumptions |
| --- | --- | --- |
| egg-incubation | 4 days or ticks | Number of days (ticks or iterations) when the ACB agent is in egg stage (dot shape) |
| larval-stage | 25 days | Number of days when the ACB agent is in larval stage (bug shape) |
| pupation | 6 days | Number of days when the ACB agent is in pupal stage (still bug shape) |
| death-age | 43 days | Number of days before ACB agent dies |
| egg-survival | 96% | Stochastically applied whether an ACB agent will survive while it is in the corresponding stage: egg, larval, or pupal stage. |
| larval-survival | 85% |  |
| pupal-survival | 90% |  |
| fitness-cost | 0.25 | Fitness cost of ACB agents who have resistant alleles (either red or violet). This is applied by multiplying to the larval survival. |
| mate-age | 36 | Number of days when the ACB turns to adult moth (butterfly shape) and starts to mate and move around |
| max-mate-count | 2 | Number of times an adult female ACB agent can mate and reproduce in the model |
| offsprings | 5 | Number of eggs that can be laid by a adult female ACB |
| mating-radius | 10 | The distance with respect to an adult female ACB agent where it can randomly choose a mate |
| max-distance | 5 | The maximum range that adult ACB agents can randomly move per iteration |
| Note: The last four parameters were calibrated due to computing power limitations. The rest of the parameters were based on the literatures for life tables and life cycles (Hussein and Ibrahim, 1992) (Caasi-Lit nd Sapin, 2012). |  |  |

We assigned two (2) genotypes of ACB: one for resistance genotype (SS, SR, or RR) used for single BT event simulations, and an additional resistance genotype (SS2, SR2, or RR2) used in combination with the first genotype for pyramid BT event simulations. With these two genotypes, we assigned three colors to ACB agents representing genotypes for both the single BT and pyramid BT events. Blue ACB agents have at least one homozygous dominant susceptible allele either SS or SS2, otherwise they will be assigned with violet or red color. Red ACB agents have RR allele for a single BT event or both RR and RR2 alleles for pyramid BT events. Violet ACB has either SR and/or SR2 heterozygous alleles and it must not have either an SS or SS2 alleles. Table 2 shows this color assignment scheme.

**Table 2.** Color assignment of ACB agents based on their genotypes.

| Color | Single BT event genotype | Pyramid BT event genotypes<br>(Single BT genotype and additional genotype2) |
| --- | --- | --- |
| **Blue** | SS | SS and SS2<br>SS and SR2<br>SS and RR2<br>SR and SS2<br>RR and SS2 |
| **Violet** | SR | SR and SR2<br>SR and RR2 |
| **Red** | RR | RR and RR2 |
| Single BT genotypes: homozygous dominant susceptible **SS**, heterozygous dominant susceptible **SR**, and homozygous recessive resistant **RR2**. Additional genotype (genotype2) for Pyramid BT: homozygous dominant susceptible **SS2**, heterozygous dominant susceptible **SR2**, and homozygous recessive resistant **RR2** |  |  |

### 2.3. Scales

Each iteration of the model is called a “tick” in NetLogo. In our model a tick is scaled to a time duration of one day. This is important to note because this model is simulated for 200 ticks, covering five generations. Each patch represents a unit area scaled to plant where the eggs are laid and where the larva will stay until pupation. Only when an ACB agent becomes an adult moth will it be able to move to other patches.

## 3. Process Overview

The methods for our ABM-IRM is summarized by the conceptual diagram presented in Figure 2. The model can be divided into two parts the initialization stage and the simulation stage. The setup or initialization stage can be further divided into the initialization of the corn field agents and the ACB agents. The simulation will mainly go through the life cycle of ACB agents.

**Figure 2.**
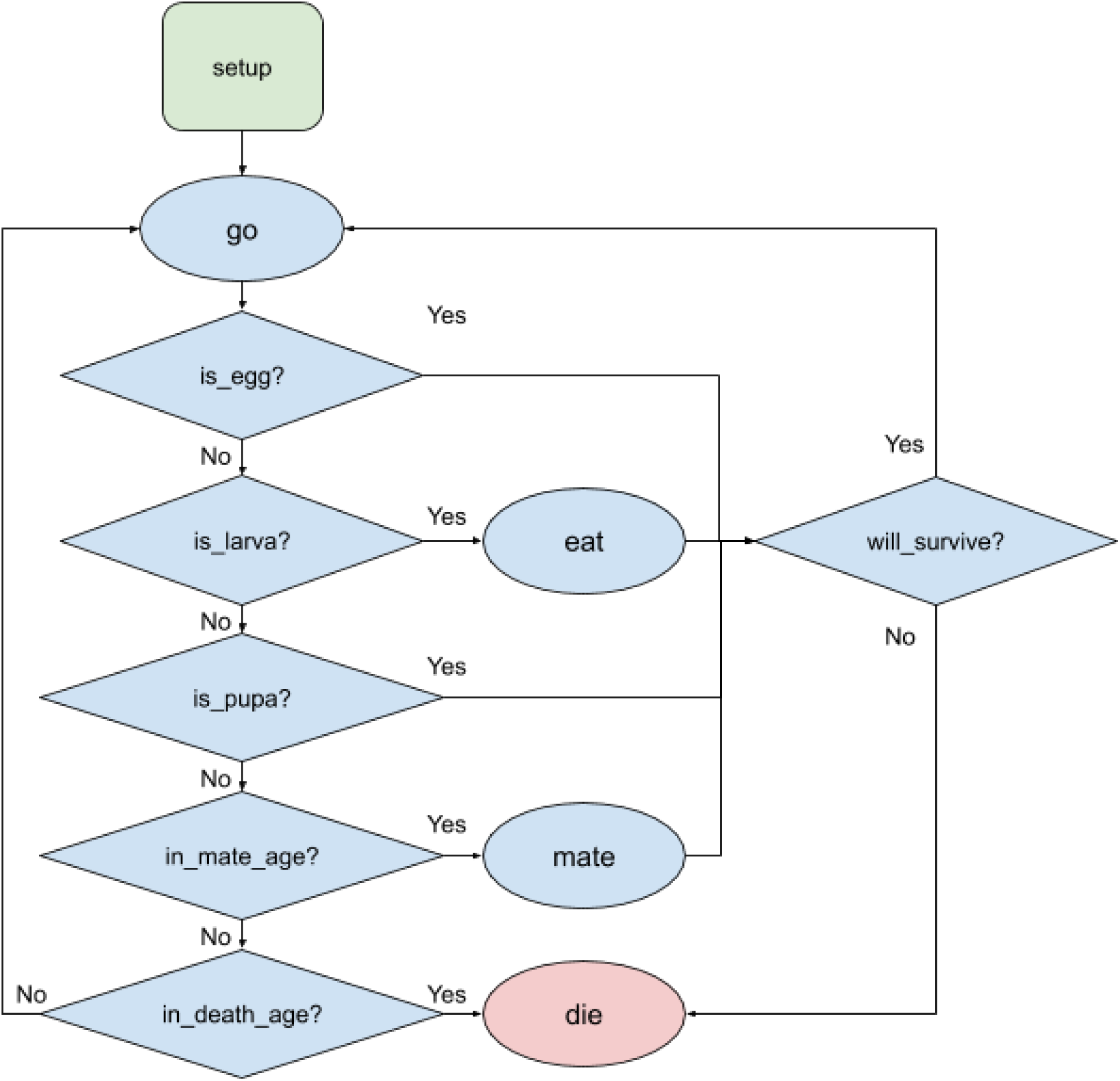
Simplified flowchart of ABM-IRM. The model starts with the setup or initialization of parameters for the refuge area and initial ACB agents. The simulation starts with the go sub-routine and loops through relevant stages of the life cycle of ACB agents.

### 3.1. Initial setup

The corn field agents are initialized by setting the percentage of refuge area and selecting the refuge type. Refuge types are shown in Figure 1. In the initial setup, 10,000 ACB agents can be created randomly with the following properties: random age from 1 - 43 days, random sex (male or female), random coordinates and stochastically assigned genotypes. The genotype of ACB agents are stochastically assigned with RR-rate usually set at 5% and SR rate at 1%, and this also the same for RR2-rate and SR2 rate, respectively. This stochastic distribution is only applied in the initial setup. The genotype of the offspring that will be created in the simulation will be solely based on the crossing of the genotypes of their parents.

### 3.2 Simulation

The simulation loops with the “go” subroutine, where agents are aged by a day for each tick or iteration. The shape and color of the agents are also updated in the “go” subroutine. For each iteration, ACB agents behave depending on their developmental stage (see Table 1). Eggs and pupa go through with the iteration checking their survival rates at 96% and 90% respectively. Interactions are mainly designed for larva and adult ACB agents.

When a larva emerges, its survival depends on its genotype and survival rates. The survival of resistant larva is checked based on the normal survival rate of ACB at 85% multiplied by 75% or (1 - fitness cost). Susceptible ACB larva dies when they emerge in a BT patch (green) and those that emerge in refuge area (yellow) are checked with normal survival rate for larva at 85%.

Only adult ACB agents can move around and mate. When an adult ACB emerges, it immediately goes through the mating process. Unlike for their life tables, the current implementation of the mating process may not really reflect the mating pattern for ACB due to computing limitations. Adult female can mate at most twice in the current version of the model. Mating starts when a viable adult female tries to find a male within its radius, set as far as ten patches. For each mating process, a female ACB can reproduce at most five (5) eggs. The eggs are hatched on the current position of their female mother, hence, the eggs may be hatched in either a refuge area or a BT patch. An adult ACB dies when it reached the death-age at 43 days. Due to computing power limitations, we terminate the simulations after 200 ticks, equivalent to 200 days or roughly five (5) generations.

## 4. Discussion of Results and Emergent Patterns

### 4.1. Baseline scenarios

When we simulate our ABM-IRM model without refuge areas, we anticipate two possible scenarios that can happen: case 1) susceptible population will be wiped out and resistant population may increase and dominate, and case 2) all the ACB population will be wiped out. We consider both cases as worst-cases in our model because the wiping out of susceptible population is contrary to the phenomenon that require significant susceptible population of ACB in refuge areas to control the resistance to BT. Snapshots of these baseline scenarios are shown in Figure 3.

**Figure 3.**
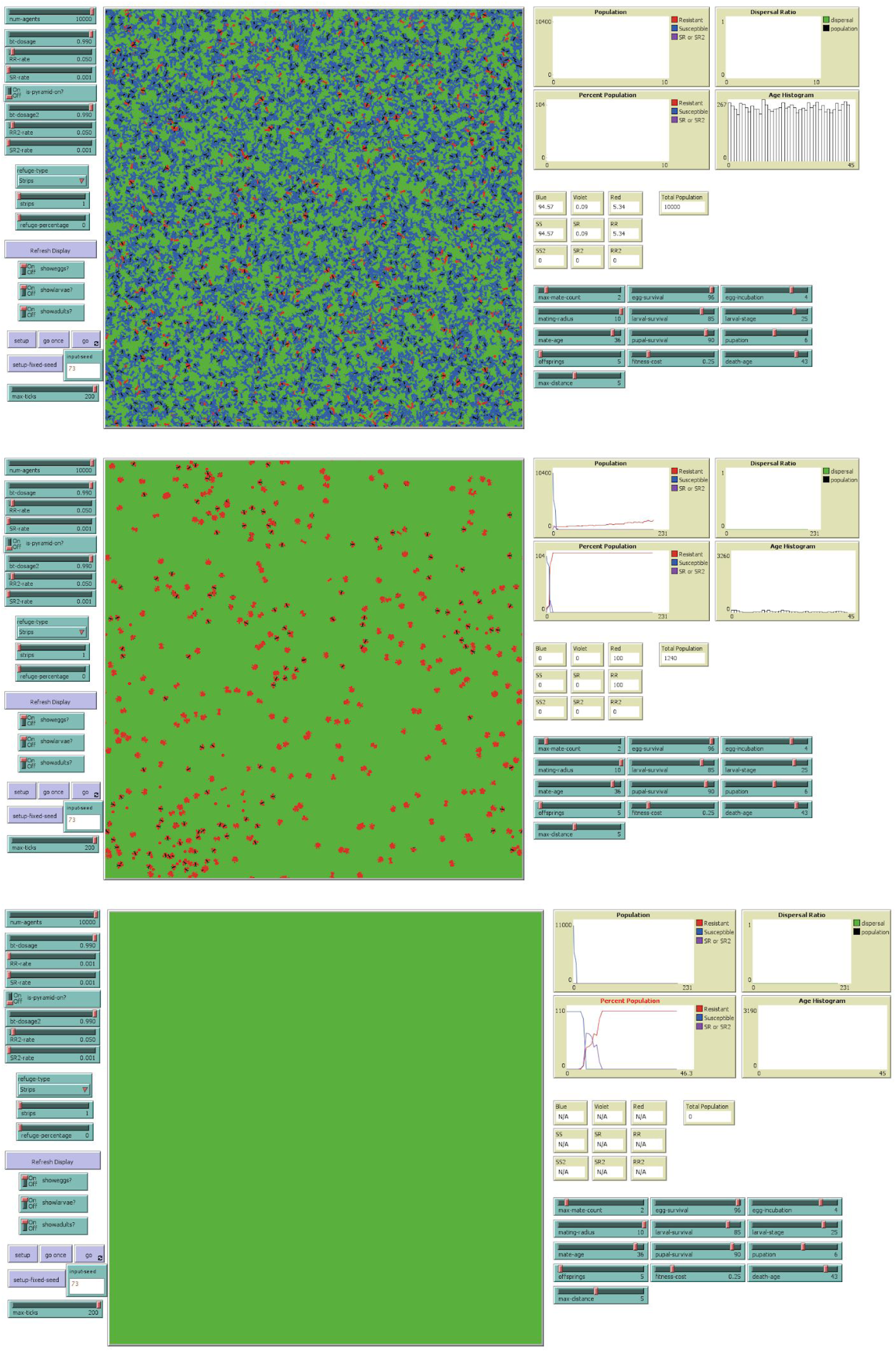
Baseline Scenarios when there are no refuge areas. The snapshot at the top is taken right after the initializing the baseline instance of the model. Middle snapshot is taken when simulation is terminated after 200 iterations. It shows susceptible population are wiped out and resistant population increased. Bottom snapshot is an instance when all ACB population are wiped out. This instance was run at 1% RR-rate instead of at 5%.

### 4.2. Single BT event at 30% and 15% refuge areas with varying refuge types

We simulated the single BT even of our model at RR-Rate at 15% and 30% refuge area percentage on the three refuge types (shown in Figure 1). Snapshots of the results of our simulations are shown in Figure 4. We can see in the snapshot for “randomseeds” refuge type, that even at higher than recommended refuge area at 30%, the susceptible population declined rapidly followed by an on set of increase of the resistant population. This is similar to Case 1 of the baseline scenario where there are no refuge area at all. However, we can see sustained increase of susceptible population for the “strip” and “circle” refuge types.

**Figure 4.**
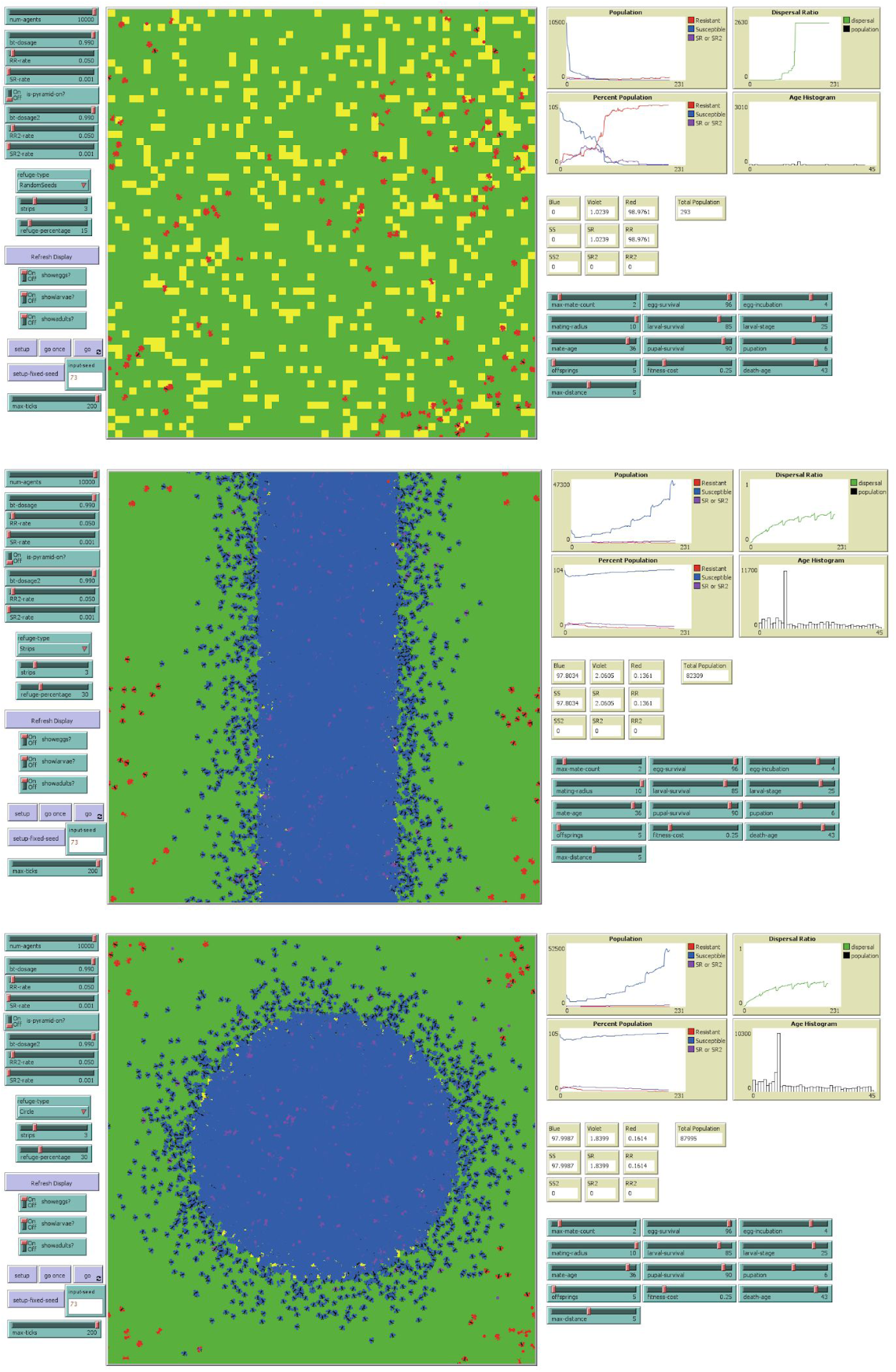
Snapshots for Single BT event with 30% refuge area and varying refuge types.

When we reduced the refuge area at 15%, maintaining RR-Rate at 5% and still in a single BT event setup, we found similar results compared to 30% refuge area simulations. However, we observed more rapid decline of susceptible population for all cases (compare Figure 4 and 5). This is an expected phenomenon when we reduced the refuge area where susceptible population can thrive. It must be noted that even in this reduced refuge area setup, “strips” and “circle” refuge types still afforded sustained susceptible population, and more effectively for “circle” refuge type.

**Figure 5.**
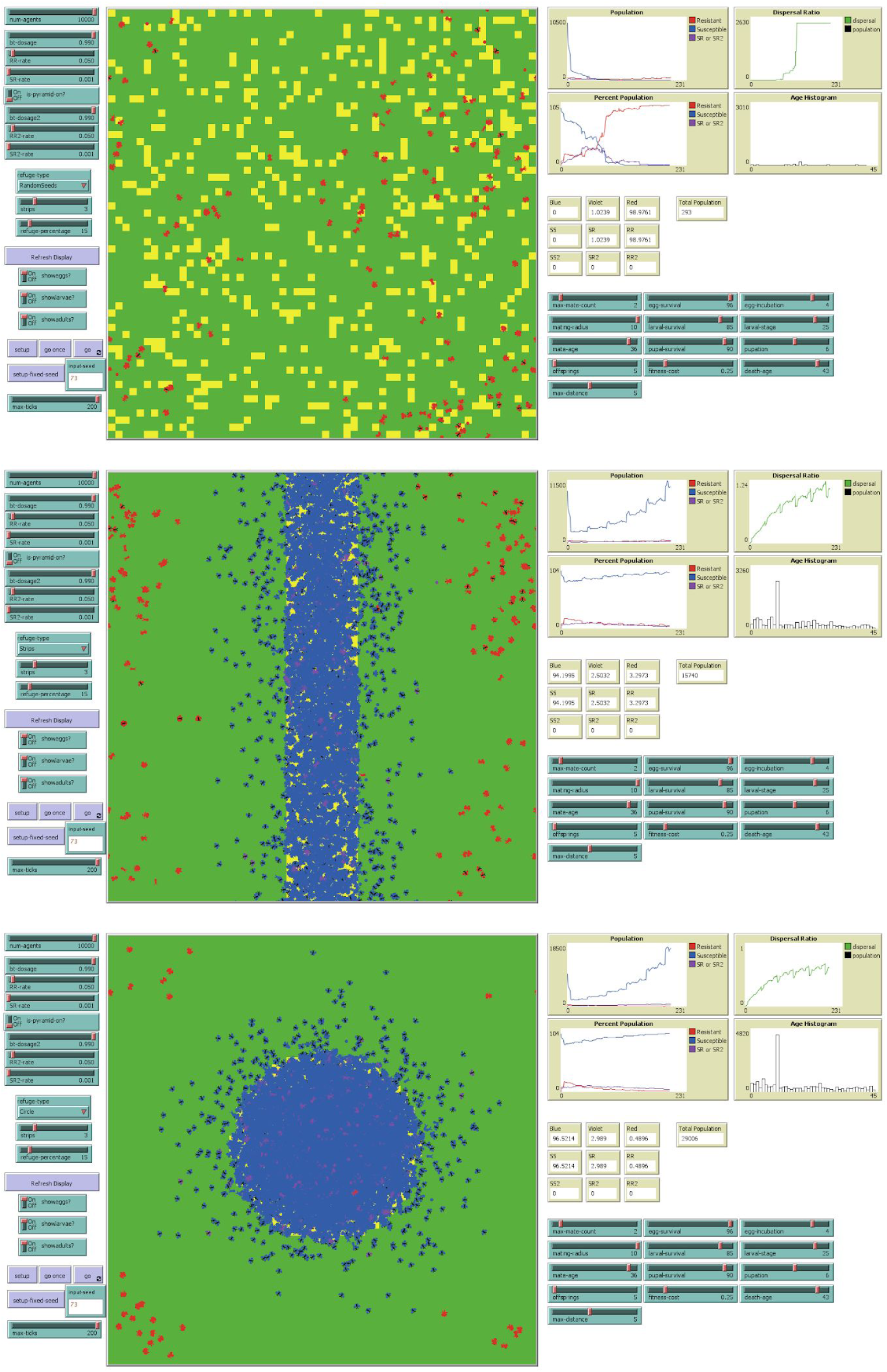
Snapshots of single BT event with 15% refuge.

### 4.3. Pyramid BT event at 15% refuge areas with varying refuge types

To test the pyramid BT event, we considered the setup with 15% refuge area (instead of 30%) and initial RR2-rate at 5% (instead of values lower than RR-rate) to highlight the difference of results for the “randomseeds” refuge type compared to the other two refuge types. In Figure 6, the snapshots of results for “randomseeds” refuge type showed that even with the reinforcement of a pyramid BT event, susceptible population still declined. In contrast, snapshots for “strip” and “circle” refuge types showed sustainability of susceptible population reinforced by pyramid BT event. Like in the single BT event simulations, “circle” refuge type afforded better sustainability of susceptible population.

### 4.4. Emergent Patterns

As shown in Figures 4,5 and 6, susceptible population decline in “randomseeds” refuge type. As discussed earlier, this may lead to the increase of resistant population. The small refuge areas generated in the “randomseeds” refuge type are randomly distributed all over the field. This can increase the dispersal of the susceptible ACB and dilute the gene pool of nearby resistant population. However,the advantage gained from this dispersal may not be able to compensate the decline of susceptible population in the small refuge areas.

We also tried simulating multiple refuge strips through the “strips” refuge type setup. We increased the number of strips for refuge areas distributed evenly, while maintaining the percentage of refuge area. When the strips became too thin, they effectively became like the small patches of “randomseed” refuge type that were not able to sustain the susceptible population. Moreover, a single strip refuge with small refuge area percentage will also not be able to sustain the susceptible population.

This led us as to define the dispersal ratio:

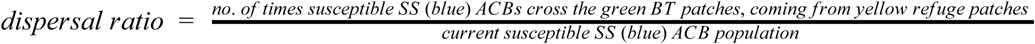

The dispersal ratio is inversely proportional to the susceptible ACB population. The slower the increase of dispersal ratio may mean increase susceptible population or susceptible ACB rapidly disperse from the refuge area and dilute the resistant population. This pattern can also be seen in the snapshots of our results presented Figures 4,5, and 6. The plots of dispersal ratio for “randomseeds” refuge type are steeper compared the plots for “strip” and “circle” refuge types. The plots for “circle” refuge type grows even slower than that of “strip” refuge type. The circle shape afforded better balance between the increase in dispersal of susceptible ACB and sustainability of the susceptible population in the refuge area, as compared to the “strip” refuge type.

### 4.5. Recommendation for the IRM of ACB to BT Corn

With the current limitations of our ABM-IRM model, i.e. calibrated mating parameters considering the computing power limitations when running our model, we could only recommend the following general ACB IRM strategies based on the emergent patterns that we have found in our model:

1. Aim for the balance of advantages between the dispersal of susceptible population in diluting the gene pool of resistant population while sustaining the susceptible population in refuge areas.
2. IRM strategies can be enforced with pyramid BT event but should be done with careful consideration to the first recommendation. For example, enforcing with “randomseeds” refuge type with pyramid BT may not be effective at all if “randomseeds” refuge type generated too small refuge areas in the first place.

---

**Figure 5.**
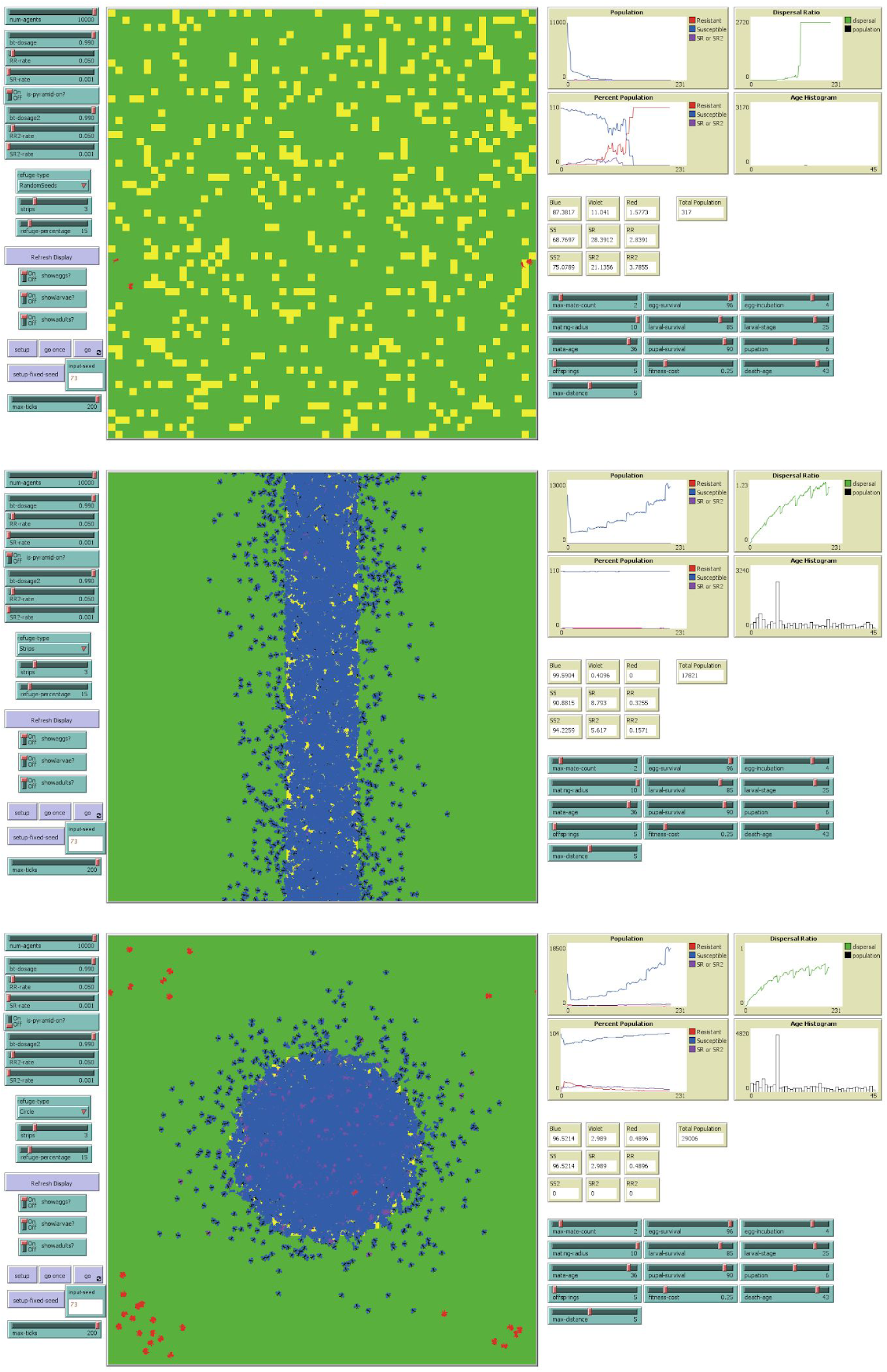
Snapshots for Pyramid BT event with 15% refuge area and varying refuge types.

